# Traces of transposable elements in genome dark matter co-opted by flowering gene regulation networks

**DOI:** 10.1101/547877

**Authors:** Agnès Baud, Mariène Wan, Danielle Nouaud, Nicolas Francillonne, Dominique Anxolabéhère, Hadi Quesneville

**Author notes:** **Cite as**: Baud, A., Wan, M., Nouaud, D., Francillonne, N., Anxolabéhère, D. and Quesneville, H. (2021). Traces of transposable elements in genome dark matter co-opted by flowering gene regulation networks. bioRxiv, 547877, ver. 6 peer-reviewed and recommended by PCI Genomics.doi: https://doi.org/10.1101/547877.

## Abstract

Transposable elements (TEs) are mobile, repetitive DNA sequences that make the largest contribution to genome bulk. They thus contribute to the so-called “dark matter of the genome”, the part of the genome in which nothing is immediately recognizable as biologically functional.

We developed a new method, based on *k-mers*, to identify degenerate TE sequences. With this new algorithm, we detect up to 10% of the *A. thaliana* genome as derived from as yet unidentified TEs, bringing the proportion of the genome known to be derived from TEs up to 50%. A significant proportion of these sequences overlapped conserved non-coding sequences identified in crucifers and rosids, and transcription factor binding sites. They are overrepresented in some gene regulation networks, such as the flowering gene network, suggesting a functional role for these sequences that have been conserved for more than 100 million years, since the spread of flowering plants in the Cretaceous.

## Introduction

Transposable elements (TEs) are mobile repetitive DNA sequences that make a major contribution to the bulk of the genome in many organisms [1–5]. They can represent up to 85% of some genomes as in wheat and maize.

TEs invade genomes, through their ability to amplify. However, they are also controlled by their host, through multiple pathways involving RNAi machinery. They invade genomes in a recurrent manner, through bursts of transposition that are rapidly halted by host defense mechanisms. TEs remain quiescent in the genome for long periods of time, until they are reactivated by events such as genomic shocks. TE sequences also accumulate mutations, which may inactivate the sequence by rendering it too degenerate to be functional. The TE sequence thus gradually “blends into” the background genome sequence until it is no longer recognizable. It then contributes to the so-called “dark matter of the genome”, the part of the genome in which nothing is immediately recognizable as biologically functional.

Little is known about the evolution and impact of TE sequences over long periods of time. We explored this question, by developing an innovative repeat annotation approach, which we call *cross-species TE annotation* because it uses closely related species to enhance detection sensitivity for ancient, degenerate repeated sequences [6]. We analyzed the genome of several relatives of *A. thaliana* that diverged about 5-40 million years (My) ago [7]. We generated a library of consensus repeat sequences that we appended to the *A. thaliana* TE reference library, to compile a “Brassicaceae” library. This compiled TE library was used to annotate the *A. thaliana* Col-0 genome to explore the effects of the long-term presence of TEs on genome evolution. Our Brassicaceae TE annotation, excluding annotations overlaping CDS, covers more than 31.8 Mb (26.7%) of the *A. thaliana* genome, and is highly sensitive for the detection of degenerate TE sequences, as it identified one third more TEs than the current official annotation [8]. The detection of many TE copies in *A. thaliana* on the basis of consensus sequences built from sequences in related species provides evidence in support of these *A. thaliana* repeats originating from the common ancestors of these species.

However, our ability to recognize the part of the dark matter derived from TEs remains limited by the sensitivity of current alignment algorithms. We present here a new tool that we developed to improve this strategy. Our new algorithm can find older and more degenerate TE sequences. Indeed, with this tool, we were able to detect up to 10% more of the *A. thaliana* genome as material derived from as yet unidentified TEs. By combining several strategies and tools, we were able to bring the proportion of the genome of this species known to be derived from TEs up to 50%. Interestingly, the new sequences detected were generally very short and located in the 500 bp immediately upstream from genes. Their epigenetic status and nucleotide composition attest to their origination from an ancient TE. Moreover, long-term conservation in orthologous positions and overlap with experimentally identified transcription factor binding sites (TFBS), suggest that they have been co-opted for new functional roles. Interestingly, these sequences were found to be overrepresented in the 5’ sequences of flowering genes. A significant proportion of these TEs overlap with TFBSs able to bind transcription factors (TFs) known to be involved in flowering. Their overlaps with conserved non-coding sequences (CNS) suggest a long-term impact of TEs on flowering, since the initial global spread of flowering plants in the Cretaceous period.

## Methods

### Genome sequences

Genome sequences were obtained from the following sources: *A. thaliana* ecotype Col-0 (TAIR10 release) (http://www.phytozome.com/arabidopsis.php); *A. lyrata* (v1.0, http://www.phytozome.com/alyrata.php); *C. rubella* (initial release, http://www.phytozome.com/capsella.php); *A. alpina* (preliminary release, courtesy of Eva-Maria Willing, George Coupland, and Korbinian Schneeberger); *Schrenkiella parvulum* (formerly *Thellungiella parvula;* v2.0, http://thellungiella.org/data/); and *B. rapa* (v1.2, http://www.phytozome.com/napacabbage.php).

### Genome annotation

TAIR10 gene and TE annotations were retrieved from the URGI website (https://urgi.versailles.inra.fr/gb2/gbrowse/tairv10_pub_TEs/).

The *Arabidopsis thaliana* “Brassicaceae*”* TE annotation was developed in a previous published study [6]. Briefly, in this previous work, for all the genomes from the five *Arabidopsis thaliana* ecotypes that have been assembled (Col-0, Ler-1, Kro-0, Bur-0 and C24), *Arabidopsis lyrata, Capsella rubella, Arabis alpina, Brassica rapa, Thellungiella salsuginea*, and *Schrenkiella parvula*, the TEdenovo pipeline from the REPET package (v2.0) [9–11] was used with default parameters and combining the similarity and structural branches. Consensus sequences derived from the structural branches, which use LTR Harvest, were retained only when they presented pfam domains typical of LTR retrotransposons. Consensus sequences were classified with REPET, by checking for characteristic structural features and similarities to known TEs from Repbase Update (17.01)[12], and by scanning against the Pfam library (26.0)[13] with HMMER3 [14]. All the consensus repeat sequences generated were compiled into a “Brassicaceae” library, which we used to annotate the Col-0 genome with TEannot from the REPET package and default settings.

### Brassicaceae TE copies

In order to annotate TE copies in each Brassicaceae genomes, we performed a new REPET analysis taking advantage of its improvements and of some new genome assembly. Hence, for all genomes from *Arabidopsis thaliana* Col-0 ecotype, *Arabidopsis lyrata, Capsella rubella, Arabis alpina, Brassica rapa* and *Schrenkiella parvula*, we used REPET package v2.5 with its two pipelines, TEdenovo and TEannot. We used the similarity branch of TEdenovo with default parameters on each genome, followed by TEannot with default parameters (sensitivity 2). From this first annotation, we selected consensus sequences containng at least one full-length copy (*i*.*e*. aligned over more than 95% of the length of the consensus sequence), on which which performed a second run of TEannot. This procedure has been shown to improve the quality of annotation [15]. Copies from the consensus annotated as ‘PotentialHostGene’ were removed.

### Prediction accuracy

True positives (TP) are defined as predicted TE nucleotides that truly belong to a TE copy. False positives (FP) are the predicted TE nucleotides that do not really belong to a TE copy. True negatives (TN) are the nucleotides correctly predicted not to belong to a TE copy (correct rejection), and false negatives (FN) are the true TE copy nucleotides missed by the TE prediction process.

Sensitivity, the true positive rate, given by the formula TP/(TP+FP), is obtained by calculating the fraction of nucleotides in the predicted TE overlapping with the TE reference annotation.

Specificity, also refered to as the true negative rate, is less straightforward to calculate. It can be calculated according to the formula TN/(TN+FP), but TN and FP are difficult to determine for TEs, as they can only be known if we are sure that we have identified all the TE copies in the genome, which does not really seem possible. However, as a first approximation, we can consider that genes are not TEs, and are not derived from TEs, and use this information to obtain more accurate estimates for TN and FP. This is obviously an approximation as TEs are known to be sometimes part of genes. Hopefully this is rare compared to other regions of the genome, in particular if we consider the coding sequence (CDS) as it discards introns as well as 5’ and 3’ UTR where TEs can be found frequent. In this context, FP are predicted TE nucleotides that overlap a gene CDS annotation, and TN are CDS regions not predicted to be TEs.

Accuracy, given by ACC=(TP+TN)/(TP+TN+FN+FP), is the rate of correct predictions.

### Epigenetic data

We used a small-RNA map from Lister et al. (2008) [16] corresponding to dataset GSM277608 from the Gene Expression Omnibus (GEO) database (http://www.ncbi.nlm.nih.gov/geo/). The original coordinates were projected onto the TAIR10 assembly. The occurrences of multiply mapping reads were distributed evenly between genomic copies. This small-RNA dataset is derived from inflorescences of plants grown at 23°C with a 16 hours / 8hours dark cycle. Small RNAs from 15 to 35 nucleotides where extracted from bulk RNA extraction.

We used the 10 chromatin mark maps (H3K18ac, H3K27me1, H3K27me3, H3K36me2, H3K36me3,H3K4me2, H3K4me3, H3K9ac, H3K9me2 et H3) from Luo *et al*. [17].

Reads overlapping an annotation were counted with the CompareOverlapping.py script (option –O) of the S-Mart package [18].

We normalized counts by calculating the ratio of the mean number of reads overlapping an annotation to the number of overlapping reads from the input.

Hierarchical clustering was performed on the epigenetic marks, based on the normalized ratio, with the *seaborn* python library, the correlation metric and a standard-scale normalization for each mark.

### TFBS and CNS data

We use TFBS data compiled by Heyndrickx *et al*. [19] from ChIP-seq experiments, which we downloaded from http://bioinformatics.psb.ugent.be/cig_data/RegNet/.

CNSs data were retrieved from the work of Van de Velde *et al*. [20], from http://bioinformatics.psb.ugent.be/cig_data/Ath_CNS/Ath_CNS.php, and that of Haudry *et al*. [21] from http://mustang.biol.mcgill.ca:8885/download/A.thaliana/AT_CNS.bed.

### Analysis of binding motifs

We searched for binding motifs with the MEME suite server [22] from http://meme-suite.org. We used MEME-ChIP [23] and JASPAR2018 CORE non-redundant databases.

### Analysis of orthologous genes

We used OrthoMCL [24] version 2.0 to identify genes orthologous between *A. thaliana, A* .*lyrata, C. rubella*, and *S. parvulum*. From the 21689 groups of orthologs obtained, we retained only 6921 for which four genes were identified, one from each species, to limit the detection of false-positive paralogs by this method.

### Statistical analysis

We used the python libraries *pynum, scipy* for statistics, *matlibplot* for graphics and *panda* for data manipulation. *Jupyter notebooks* were used to monitor the analysis.

### Sequence and coordinate manipulation

We obtained random sequences with *shuffle* from the SQUID 1.9g package [25] and *revseq* from the Emboss 6.1.0 [26] package.

Genome coordinates were manipulated with the S-Mart package [18]. In particular, we used modifyGenomicCoordinates (version 1.0.1) and CompareOverlapping (version 1.0.4) to extend coordinates within the 5’ region of genes, and find overlaps, respectively.

## Results

### Duster: a new approach for analyzing old degenerate transposable elements

Following their divergence from a common ancestor, repeat families have different destinies in different genomes. A repeat family may stop multiplying in one species, but may continue to multiply in another closely related species. The burst of transposition in an autonomous repeat family is a highly selective process: only the copies that have accumulated limited numbers of mutations remain functional and are able to transpose during the burst. This selective burst of transposition drives multiplication of the best conserved copies, *i*.*e*. those most similar to the ancestral sequence. Therefore, the TE families that remain active in some genomes should more similar to the ancestral sequence for longer than the decaying pool of related sequences in other genomes. Consequently, a repeat copy from one species may be considered to be relatively old if it closely resembles a sequence obtained from another species.

We previously showed [6] that identifying TEs in a species by comparison with reference sequences found in the studied species but also in closely related species leads to the detection of older TE copies than searches exclusively with the reference sequence from the study species. Indeed, this approach detects old TE sequences that would not otherwise be recognized.

We developed a program called Duster that compares a genome sequence, here considered as a query sequence, to a large number of TE sequences, *i*.*e*. a sequence library. Its algorithm used *k-mers* of size *w* (parameter *-w*) to search for similar sequences without the need to generate nucleotide aligments (figure 1). Hashed *k-mer* values can be used to speed up the search. Sensitivity is achieved by allowing one mismatch in *k-*mers every *n* consecutive nucleotides (parameter *-k*). Details of the algorithm are provided below, but it can be summarized as comparing *k-*mers between the genome shifting on the genome by few nucleotides (parameter *-S*), and each sequence from the library shifting by 1 nucleotide, and reporting matches when at least two *k-*mers are found on the same alignment diagonal (i.e. the differences between the coordinates in the query and the sequence library are identical) with a maximal distance of *d k-*mers (parameter *-d*). The region bounded by the two-extreme *k-*mer position are reported as matching. Two matching regions on the genome separated by less than *x k-*mers are merged (parameter *-f*). At the end of this first pass, the region identified on the genome can be used as a new sequence library for a new search (the *-n* parameter). This procedure is repeated until genome coverage increases by less than 1% if *-n* is set to 0.

**Figure 1:**
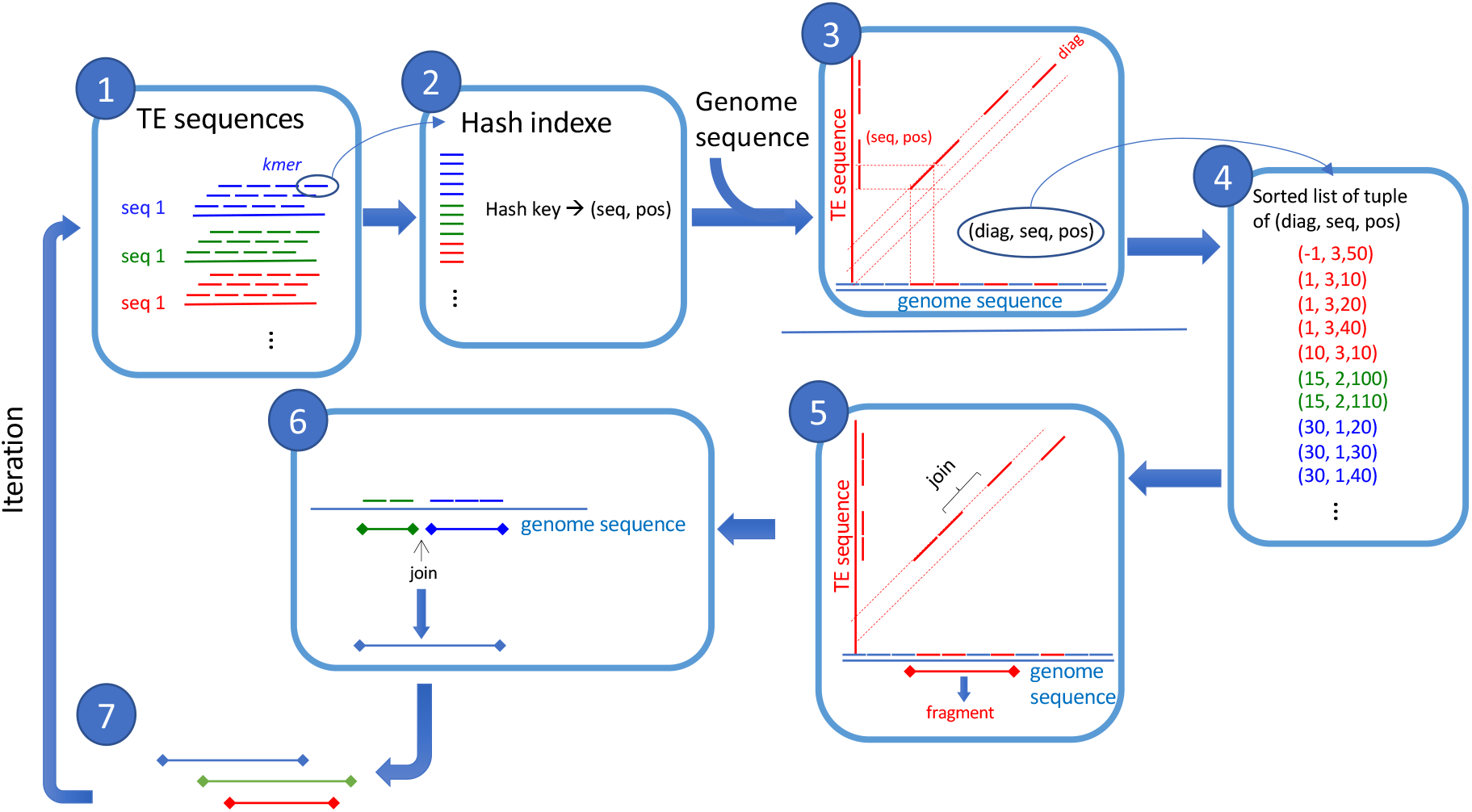
Duster algorithm overview. 1-TE sequences are cut in overlapping kmer of length w. 2-Build a hash table from kmer hash value to store their positions and sequence numbers. 3-Search kmer on genome sequence that match a TE sequence kmer, record diagonal (difference of start sequence position) with TE sequence number and position on TE sequence. 4-Sort the list of tuple by TE sequence numbers, diagonals, and positions. 5-Find consecutive kmer matches in the sorted list and join if they are on the same TE sequence, same diagonal, and with position distant by less than d kmer. Obtain fragments. 6-Join consecutive fragments on the genome sequence if distant by less than f nuclotides. 7-Take the fragment sequences as a new TE sequence set for another algorithm iteration.

### The Duster algorithm

#### Definition and notations

We consider the problem of searching for occurrences of part of a query sequence *Q* within a library of subject sequences *L = {S*_*1*_, *S*_*2*_, …, *S*_*d*_*}*. Each sequence in L is labeled with an index value *s*. We use the term *k-*mer to denote a contiguous sequence of DNA bases that is *w* bases long. The offset of a *k-*mer within a sequence *S* is the position of its first base with respect to the first base of *S*. We use the letter *j* to denote offsets and the notation *S*_*j*_ to denote the *k-*mer of *S* that has an offset *j*. The position within *L* of each occurrence of each *k-*mer may then be described by an *(s, j)* pair.

#### *k*-mer hash function

We store *k-mer* as a *hash-value* to speed up comparisons and to reduce memory requirement. The hash-value is obtained from the *k-*mer nucleotide sequence, by encoding each of the four possible nucleotides as two binary digits, as follows: f(A or any value different from C, G, T) = 00_2_, f(C) = 01_2_, f(G) = 10_2_, f(T)= 11_2_. With this encoding system, any *k-*mer *K = b*_*1*_*b*_*2*_ … *b*_*k*_ can be represented in a unique manner by a *2k* bit integer

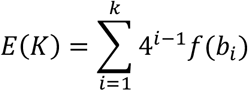

#### Constructing the hash table

The first stage of the algorithm is the conversion of *L* into a hash table. The hash table is stored in memory as two data structures, a vector of positions *P* and a vector *V* of pointers into *P*. There are *4*^*k*^ pointers in *V*, one for each of the *4*^*k*^ possible *k-*mers. The pointer at position *E(K)* of *V* points to the entry of *P* describing the position of the first occurrence of the *k-*mer in the library *L*. We can obtain the positions of all occurrences of *K* in *L* by moving *P* along the sequence from this point until it reaches the location indicated by the pointer at position *E(K)*+*1* of *V*. The hash table is constructed by making two passes through *L*. On each pass, we consider only non-overlapping *k-*mers in *S*. With the first pass, we count all nonoverlapping occurrences in *L* of each possible *k-*mer. We then use these data to calculate the pointer positions for *V* and allocate the correct amount of memory for *P*. We ignore all *k-*mers with a frequency of occurrence exceeding a cutoff threshold. This has the advantage of both reducing the size of the hash table and effectively filtering out spurious matches from low-complexity DNA sequences (see below). We make a second pass through the data, using *V* to place the *k-*mer positions in *P* at the correct positions. During these two passes, any ambiguous or unrecognized characters, such as “N”, are translated into “A”. Stretches of unrecognized characters are thus translated into *k-*mers containing only “A”s. This is generally the commonest *k-*mer in the genome and will therefore be excluded from matching by the cutoff threshold.

#### Sequence search

We use the hash table to search for occurrences of a query sequence *Q* within *L*. We proceed along *Q*, from base 0 to base *n-w*, where *n* is the length of *Q*. At any base position *t*, we obtain the list of *r* positions of the occurrences of the *k-*mer *Q*_*t*_. These positions are given by *E(Q*_*t*_) in *V*. We take this list of positions:

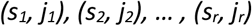

and use it to compute a list of hits:

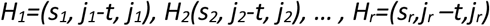

We add these hits to a cumulative list *M* of hits that grows as *t* moves from 0 to *n-w*. Depending on the sensitivity required, we shift *t* by a few nucleotides. The elements of a hit are referred to, from left to right, as the *index, shift*, and *offset*. At the end of the sequence *Q, M* is sorted, first by *index* and then by *shift*. The last step is to scan *M*, looking for consecutive hits for which the *index* and *shift* are identical. Two hits are considered consecutive if they are separated by less than *d k-*mers. Note that such hits correspond to a succession of *w* bases consistent with an alignment between *Q* and a particular sequence *S* from *L*. If we sort these hits by *offset*, we obtain regions of exact matching between the two sequences. We retain only matching regions with at least two consecutive *k-*mers separated by fewer than *d* non-overlapping *k-*mers. We can create gapped alignments including mismatches, by joining exact matching regions if they are sufficiently close on the genome sequence (parameter *-f*). The method described above finds matches only on the direct DNA strand. For the identification of matches on the reverse strand, the search is simply repeated on the sequence complementary to *Q*.

#### Improving sensitivity

In the algorithm described above, hits are obtained only if two sequences contain identical *k-*mers, limiting the sensitivity of the search. Indeed, if two sequences have diverged considerably, it is likely to be difficult to find exactly matching *k-*mers. We therefore introduced some flexibility into the *k-*mer matching procedure, by modulating the hashing function. When calculating the *k-*mer hash value, we skipped one in every *n* nucleotides. Hence, two *k-mers* with different bases may have the same hash values if the difference concerns the skipped nucleotides. If they have the same hash value, they are considered to match according to the algorithm.

#### Filtering options

*K-*mers can be removed from the initial hash table if they fail to fulfill certain criteria. First, we remove all *k-*mers containing only ‘A’, as such *k*-*mers* are overrepresented in most genomes. Note that in masked sequences, ‘N’ and ‘X’ are also converted to ‘A’ in our hashing function, automatically removing masked stretches of a sequence. For each *k-mer*, we also calculate the ratio of observed to expected occurrences, and remove all occurrences with a ratio below 1, which can be explained by chance alone. The expectation is calculated with a probabilistic background, modeled by a Markov chain in which order is a parameter (order 1 by default). Some *k-*mers can be removed, according to a threshold set by the user. Thus, *k-mers* for which the ratio of observed to expected occurrence is below a given threshold (1 in the example above) can be removed. This ratio shows to what extent the observed number of occurrences diverges from expectations. Overrepresented *k-mers* are removed as they are considered not to be specific enough. We remove those with occurrences above a given percentile (with 100% as the default setting). *K-*mers can also be removed on the basis of high entropy and low diversity, calculated as the number of different *k-*mers of the size of the background Markov chain order, divided by the maximum number of *k*-mers of this size.

#### Iterative search

The sequences of the library *L* may be considered as a sample of sequences to be searched for the sequence *Q*. Consequently, they may not contain all the *k-*mers required for a sensitive detection of *Q* in *L*. We thus implemented an iterative search to enrich the *k-mer* repertoire with the sequence *Q*. We reran the algorithm described above, replacing *L* with the regions identified in *Q*, and filtering out sequences below a threshold length. This procedure can be repeated until *Q* coverage changes by less than 1% between successive iterations.

### Assessment of the performance of Duster

We assessed the performance of Duster, by calculating its prediction accuracy. This accuracy (ACC) was obtained by calculating the sensitivity (Sn) and specificity (Sp) of the predicted TE annotation by comparing the prediction with a reference annotation at nucleotide level (see Materials and Methods). We used the official annotation for *A. thaliana* from TAIR as the reference here. ACC takes both Sn and Sp into account, providing a convenient aggregate value. We therefore decided to maximize this value in our benchmark tests, for which the sequence library used was that of TE sequences from the TAIR annotation.

We chose the parameter set that gave the best results for Duster in our hands empirically, by optimizing annotation accuracy with TE copy sequences from other Brassicaceae species (data not shown). We used this parameter to compare Duster performances, benchmarking with tools implementing other algorithms that could be used for similar analyses. For this comparison, we chose BLAST [26] and MegaBLAST [27], two widely used sequence comparison algorithms. Neither of these tools was designed to be run on a long genomic sequence. We therefore ran them through Blaster [28], which pre- and post-processes the input sequence and output results, respectively, to facilitate their use.

Table 1 shows the results obtained with Duster, BLAST and MegaBLAST. Duster was run twice, varying the distance required between two *k-*mers (*–d* parameter), using the values of 0 or 5, and the position shift on the genomic sequence (*-S* parameter), using values of 15 (size of the *k-*mer) or 7 (*k-*mers overlapping by 7 bp). The parameter *(–d 0; –S 15*) is, by definition, less sensitive than (*-d 5; -S 7*). Note that parameter *–f* is the maximum sequence distance beween two consecutive fragments on the genome sequence to be joined, and *–n* is the number of iterations. Here, the two sets of parameters are set to consider only sequences of more than 100 bp in length, with only one iteration. We also chose a *k-*mer length of 15 (parameter *–w*) and a potential nucleotide mismatch every four nucleotides (parameter *–k*).

**Table 1:** Comparison of tool performances with TE sequences from the official TAIR A. thaliana TE annotation. *computed on a Linux workstation with Intel Xeon® CPU E3-1270 v3 @ 3.50 GHz x 8 and 15.5 Go of RAM.

| Tool | Parameters | Cov. | Sn | Sp | Time* |
| --- | --- | --- | --- | --- | --- |
| Duster | -w15; -k4; -d0;<br>-f100; -S15; -n1 | 0.27<br>(shuff.: 0%<br>rev.:0.1%) | 0.99 | 0.91 | 5.0 m |
| Duster | -w15; -k4; -d5;<br>-f100; -S7; -n1 | 0.31<br>(shuff.: 0.1%<br>rev.:1%) | 0.99 | 0.87 | 7.0 m |
| Blaster/MegaBLAST | -S2 ; -L200 | 0.20 | 0.96 | 0.99 | 1.18 h |
| Blaster/MegaBLAST<br>4 Threads | -S2 ; -L200 | 0.20 | 0.96 | 0.99 | 38 m |
| Blaster/BLAST | -S2 | 0.23 | 0.99 | 0.98 | 17 h |
| Blaster/BLAST<br>2 Threads | -S2 ; -L200 | 0.22 | 0.97 | 0.98 | 8.8 h |
| Blaster/BLAST<br>4 Threads | -S2 ; -L200 | 0.22 | 0.97 | 0.98 | 6.15 h |

We show that Duster outperforms the other tools in terms of speed, taking only five to seven minutes, versus 38 minutes at best with MegaBLAST run in parallel on four threads. Sensitivity was higher for Duster and BLAST, at 0.99. Duster had a lower specificity, but a higher coverage, suggesting that our tool detects many more previously unidentified potential TEs. As a different way of assessing the false-positive rate, we ran Duster on a shuffled genome sequence respecting dinucleotide composition, and a reversed but noncomplementary sequence. Coverage remained below 0.001 for the shuffled sequence, and below 0.01 for the reversed sequence.

Based on the way we compute false positives, this suggests that many genes have regions derived from old TEs not detected with other tools. As this is one purpose of our new tool, we consider this to be a positive result, particularly given that it is not biologically inconsistent.

### Transposable elements account for up to 50% of the A. thaliana genome

Assuming that Duster would be able to detect interesting new TE sequences in the *A. thaliana* genome, we ran an analysis with all the Brassicaceae TE copies from *Arabidopsis thaliana, Arabidopsis lyrata, Capsella rubella, Schrenkiella parvulum, Arabis alpina*, and *Brassica rapa* that we had previously annotated with the REPET package (see Material and Methods). We used the parameter setting with *–d* 5 and *–S* 7, but changed *– n* to 0, allowing iteration of the algorithm until it reached a genome coverage difference between two successive iterations of less than 1%.

The TAIR10, Brassicaceae and Duster TE annotations together accounted for 49.75% of the genome sequence. This figure is 29.72% higher than that for the TAIR10 TE reference annotation (20.03%), and 10.60% higher than that for the Brassicaceae TE annotation (39.15%).

### Structural properties of Duster-specific copies

We characterized the new set of repeats identified by Duster, by using the annotations to extract copies that did not overlap with any Gene, TAIR10 TE, Brassicaceae, or *A. thaliana* REPET annotations (see Materials and Methods). We identified 19608 TE copies that were Duster-specific. We did the same for the TAIR10 and Brassicaceae annotations, thereby obtaining 177 TAIR10-specific and 5139 Brassicaceae-specific copies, by removing any copies with no overlap to another annotation.

We characterized these copies by comparing their length, chromosome distribution, and position relative to genes (figure 2). Duster-specific copies appeared to be significantly shorter than Brassicaceae-specific, TAIR10-specific, and TAIR10 copies (Figure 2A, chi-squared *p-*value respectively 3.09 × 10^−192^, 2.70 × 10^−8^, <10^− 293^). Figure 2B shows the distance to the closest 5’ or 3’ TE copy for each annotated gene. Duster-specific copies are more abundant close to genes than other copies (all chi-squared *p-*value <10^−293^, versus *Brassicaceae*-specific, TAIR10-specific, and TAIR10 copies). Similarly, Brassicaceae-specific copies were more abundant than TAIR10-specific copies. They were more frequently found upstream from genes (Figure 1B, chi-squared *p*-value <10^−293^), as were Brassicaceae-specific, and TAIR10 TE copies (all chi-squared *p*-value <10^−293^). Figure 2C shows the distribution of TE copies over the chromosomes. It shows that Duster TE copies, and, to a lesser extent, Brassicaceae TE copies, follow the chromosomal distribution of genes (see the right panel of figure 2C), whereas TAIR10 TEs follow the opposite pattern. Duster and Brassicaceae TEs have a different chromosomal distribution from the annotated TEs from TAIR10.

**Figure 2:**
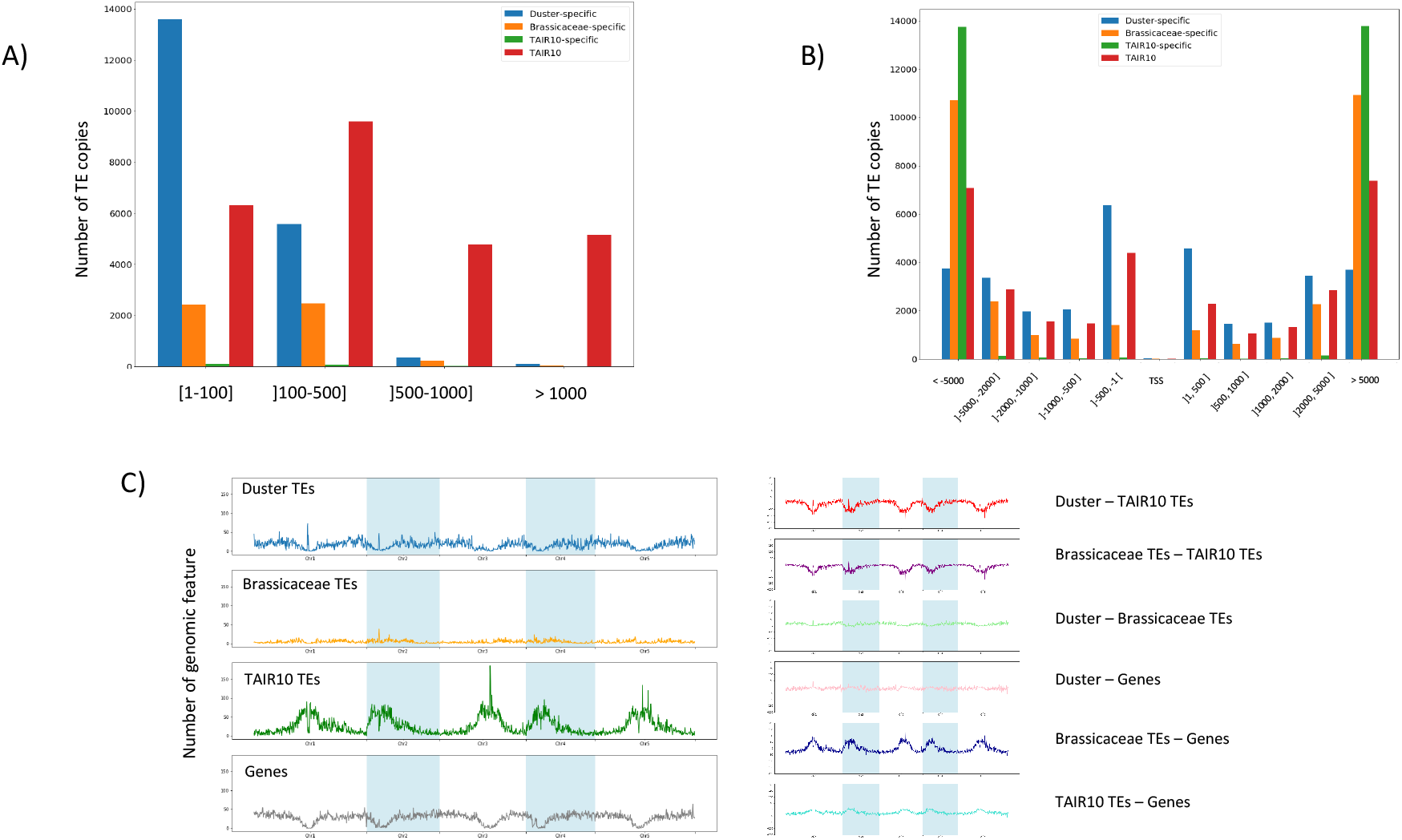
Structural characteristics of Duster-specific, Brassicaceae-specific, TAIR10-specific, and TAIR10TE copies. (A) TE length distribution, (B) distance to the closest 5’, or 3’ TE copy for each annotated gene, (C) TE copy distribution on the chromosomes. Left panel: density plot, for 100 kb windows overlapping by 10 kb. Right panel: density differences, in 100 kb bins.

Finally, we investigated the nucleotide composition of the sequences, including dinucleotides. The counts are presented as a radar plot in Figure 3A. The profile is similar for all TE copies (Duster-specific, Brassicaceae-specific, TAIR10-specific, and TAIR10 TEs). Interestingly, TAIR10-specific copies had the strongest bias towards TT, AA, AT, and TA dinucleotides, followed by Duster-specific copies. These biases, also shared by other TE copies but to a lesser extent, were thought to be a consequence of the process by which methylated cytosine is deaminated. The greater “A-T” richness of TAIR10-specific and Duster-specific copies compared to TAIR10 TE copies may indicate that they have undergone a mutation over a longer period and are therefore more ancient.

**Figure 3:**
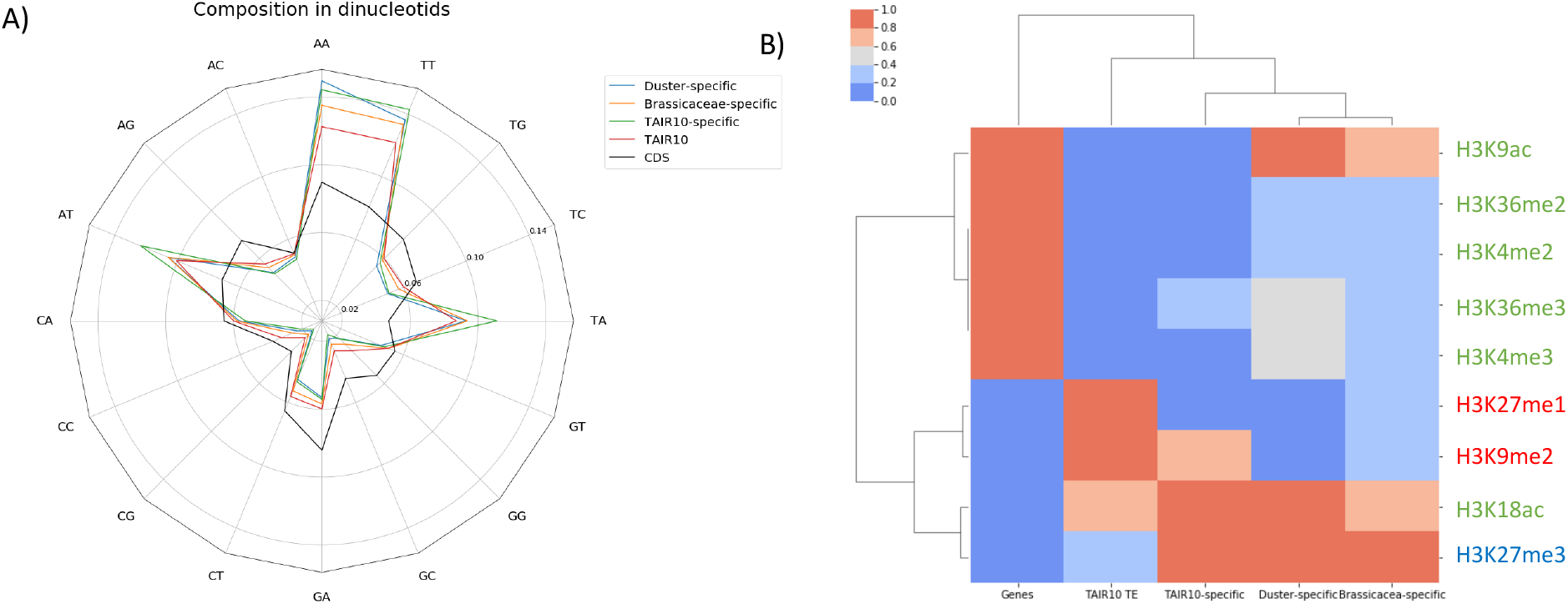
Composition of Duster-specific, Brassicaceae-specific, and TAIR10-specific copies. (A) Radar plot of the dinucleotide composition of the sequences. (B) Hierarchical clustering of TEs and genes with respect to heterochromatin marks (H3K27me1 and H3K9me2) and euchromatin marks (H3K18ac, H3K27me3, H3K36me2, H3K36me3, H3K4me2, H3K4me3, and H3K9ac). Histone marks colors denote the the type of chromatine. Green marks characterize transcriptionally active genes. Blue marks pinpoint genes transcriptionally repressed by transcriptional inhibitors. Red marks identify closed chromatin, also called heterochromatin.

### Epigenetic profiles

We investigated the epigenetic status of the identified TE copies, considering small RNAs, and chromatin marks. The small RNAs which sizes range from 15 to 35 nucleotids were taken from Lister et al. [16], for which mapped data were available. There were 4.17%, 20.14%, 16.95%, and 60.44% of matching TE copies from the Duster-specific, Brassicaceae-specific, TAIR10-specific, and TAIR10 TE datasets, respectively, in the intersection between this dataset and our annotations, indicating a low targeting by small RNA of the method-specific annotation compared to TAIR10 annotated TEs.

We analyzed nine epigenetic marks from Luo *et al*. [17], also available as mapped data. The hierarchical clustering algorithm identified distinctly different profiles for genes and for TAIR10 TEs (figure 3B). TAIR10 TEs were enriched in heterochromatin marks H3K27me1 and H3K9me2, and genes with euchromatin marks H3K36me2, H3K36me3, H3K4me2, H3K4me3, and H3K9ac as expected.

The clustering algorithm associated Duster-specific, Brassicaceae-specific, and TAIR10-specific copies with the TAIR10 TE profile, indicating that their profiles were more similar to a typical TE profile than to a gene profile. However, Brassicaceae-specific, TAIR10-specific, and Duster-specific marks copies that had very similar profiles which differ from TEs. Their copies appeared to have very few heterochromatic, however the euchromatin marks H3K27me3 and H3K18Ac are predominant for method-specific TEs. Interestingly, H3K27me3 is known to be a repressive mark preferentially associated with genes expressed at low levels or in a tissue-specific manner [29–32] and H3K18ac is usually associated with promotors [33].

### TE conservation in flowering plants

We investigated the conservation of TE copies by searching for overlaps with known conserved non-coding sequences (CNSs) identified in previous studies. We compared the TE copies with CNSs identified in crucifers [21] and rosids [20]. For both datasets, a substantial proportion of the TE copies overlapped with these CNSs (5.32 to 19.8%, see table 2). Some TE sequences overlapped with CNSs conserved in 12 rosid species: *Arabidopsis thaliana, Carica papaya, Glycine max, Malus domestica, Populustrichocarpa, Fragariavesca, Medicago truncatula, Lotus japonicus, Theobroma cacao, Ricinus communis, Manihot esculenta*, and *Vitis vinifera*. Fossil rosids dating back to the Cretaceous period, estimated by a molecular clock between 125 and 99.6 million years ago [34–36], have been found. Our findings therefore reveal a remarkable conservation of 1521 and 1213 TE insertions identified by the Duster and Brassicaceae methods, respectively, over more than 100 million years, twice as many as can be detected with the traditional annotation approach available for the TAIR10 TE annotation. We also show here that the Duster approach can detect more TEs overlapping with CNSs than the Brassicaceae method.

**Table 2:** Number of CNSs and percentage overlap with TE annotations

|  | Crucifer CNSs | Rosid CNSs | Rosid CNSs – score 11 |
| --- | --- | --- | --- |
| **All Duster** | 14774<br>14.1% | 20758<br>19.8% | 1521<br>1.45% |
| **All Brassicaceae** | 7798<br>9.39% | 13702<br>16.5% | 1213<br>1.46% |
| **All TAIR10 TEs** | 1583<br>5.32% | 4934<br>16.6% | 527<br>1.77% |
| **Duster-specific** | 3659<br>18.7% | 2580<br>13.2% | 194<br>0.99% |
| **Brassicaceae-specific** | 1848<br>36.0% | 2535<br>49.3% | 246<br>4.79% |
| **TAIR10-specific** | 27<br>15.2% | 27<br>15.2% | 3<br>1.69% |
| **5' Duster-specific** | 745<br>15.5% | 968<br>20.2% | 65<br>1.36% |
| **5' Brassicaceae-specific** | 270<br>30.6% | 452<br>51.3% | 26<br>2.95% |
| **5' TAIR10-specific** | 2<br>6.67% | 4<br>13.3% | 0<br>0.0% |

The 65 rosid CNSs from the 12 rosid species associated with Duster-specific copies in the 500 bp upstream of genes included 58 Duster-specific sequences. A MEME-ChIP [22] analysis of these sequences identified a significant 15-nucleotide TTTTTTTTT(G/T)TTT(G/T)(G/C) motif (E-value 3.4 × 10^−6^) in 27 sites. This motif matched MA1281.1 (AT5G02460), MA1274.1 (OBP3), MA1278.1 (OBP1) MA1268.1 (AT1G69570), MA1267.1 (AT5G66940), MA1272.1 (AT2G28810), MA1371.1 (IDD4), MA1279.1 (COG1), MA1156.1 (JKD), MA1374.1 (IDD7), MA1160.1 (AT1G14580), MA1158.1 (MGP), MA1157.1 (NUC), MA1125.1 (ZNF384), MA1159.1 (SGR5), MA1277.1 (Adof1), and MA1370.1 (IDD5) with all q-values < 5 × 10^−2^, described in the JASPAR database [36] as C2H2 zinc finger factors of the Dof-type. Interestingly, some of these motifs were found to be related to: (i) AT3G55370 (OBP3), which is known to encode a nuclear DOF domain-containing TF expressed primarly in roots that is responsive to salicylic acid in leaves and petals, (ii) AT1G69570 (CDF5), which is invoved in flower development and photo-periodism, (iii) AT1G29160 (COG1), which acts as a negative regulator in phytochrome-mediated light responses, (iv) AT3G50410 (OBP1), which acts as a positive regulation of transcription and play an important roles in plant growth and development and (v) AT5G02460.1, which is probably involved in early processes in vascular development.

We then looked in detail at the conservation of Duster-specific, Brassicaceae-specific, and TAIR10-specific copies in the Brassicaceae. We considered only regions close to orthologous genes found with OrthoMCL (see Materials and Methods). We focused on *A. thaliana, A. lyrata, C. rubella*, and *S. parvulum*, as these species have divergence times of 5 to 40 My. Overall, we found 6265 groups of four orthologous genes (one gene for each species) containing a TE copy annotated (available in one of the TAIR10 TEs, Brassicaceae, or Duster datasets) within the gene extended by 500 bp upstream (29% of *A. thaliana* genes). Regions encompassing both the orthologous gene and its upstream sequence were pairwise aligned with the cognate *A. thaliana* region. Within this 6265 regions, we found 2457, 353, and 11 TE copies from respectively Duster-specific, Brassicaceae-specific, and TAIR10-specific datasets. We considered a TE copy to be present if more than 50% of the *A. thaliana* annotated TE copy nucleotides were identical in the pairwise alignment. The TEs were oldest in the Duster-specific set, followed by the Brassicaceae-specific set, as shown by the height of the “111” bars of the histogram, which corresponds to the presence of a TE at orthologous positions in all four species (Figure 4). Interestingly, the “000” bars was also quite high. This bar corresponds to TEs found only in *A. thaliana*, but which belonged to method-specific sets and therefore escaped the reference *A. thaliana* TE annotation or TE detection by the simple REPET *de novo* procedure limited to *A. thaliana*. They were therefore detectable only with TEs found in other species. These copies may result from horizontal transfer from these other species, or may simply have been identified in other genomes because they are better conserved in those genomes or have enough copies to built reliable consensus. Consequently, we probably identified here TEs that had a poor success of invasion following the horizontal transfer as there are too few copies in each family, to caracterise them properly. This result illustrates the utility of our cross-species TE annotation approach and the greater efficiency of Duster than of the REPET annotation procedure.

**Figure 4:**
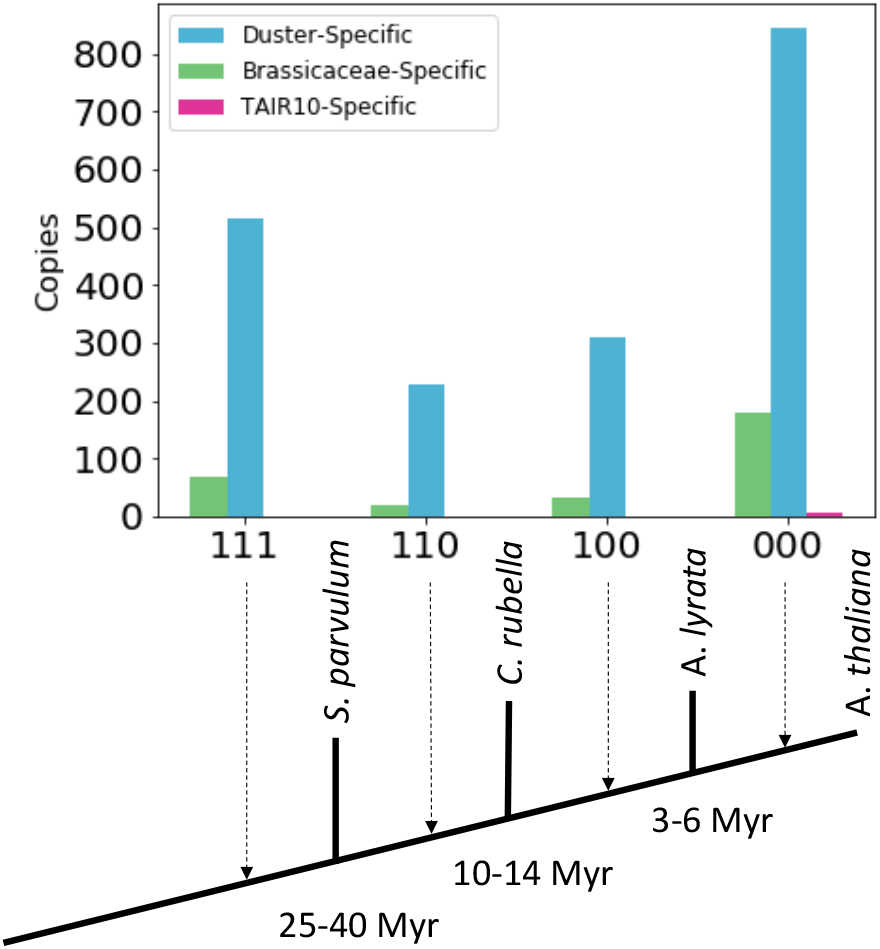
Numbers of sequences conserved in orthologous positions between species. The three-digit code indicates the species in which the sequences were present. A “1” indicates presence and a “0” absence, in A. lyrate (position 1), C. rubella (position 2), and S. parvulum (position 3).

### Contribution of TEs to the architecture of gene regulatory networks

We investigated the functional role of these TE sequences, which may have been co-opted for some regulatory purpose. We chose to study two gene regulatory networks (GRNs) in which TEs might be suspected to play a role. The genes controlling flower development in *Arabidopsis thaliana* are good candidates, as some alleles have been reported to be controlled by a TE sequence in *A. thaliana*: the FLOWERING WAGENINGEN (FWA) locus [37], and FLOWERING LOCUS C (FLC) [38,39].

We considered the genes reported by Chen et al. 2018 [40] in their paper describing the architecture of GRNs controlling flower development in *Arabidopsis thaliana*. We searched the 500 bp immediately upstream from these genes for the presence of Duster-specific, Brassicaceae-specific, and TAIR10-specific TEs.

An enrichment in Duster-specific regions was observed in the 5’ sequences of flowering genes: 33.1% of these sequences contained such regions, versus only 17.5% for all genes (chi-squared *p*-value=2.4 × 10^−7^, table 3). Brassicaceae-specific regions and specific regions identified by TAIR10 displayed no particular enrichment, with 4.46% versus 3.21% (chi-squared *p*-value=0.37), and 0% *vs* 0.11% (chi-squared *p*-value=0.68), respectively of 5’ sequences containing such regions.

**Table 3:** Method-specific TEs overlap with the 500 bp immediately upstream from genes

|  | Duster-specific | Brassicaceae-specific | TAIR10-specific |
| --- | --- | --- | --- |
| All genes | 17.5% | 3.21% | 0.11% |
| All flowering genes | 33.1% | 4.46% | 0.0% |
| Stress genes | 16.5% | 4.71% | 0.0% |

We futher explored the overrepresentation of Duster-specific and Brassicaceae-specific TEs in GRNs, by focusing on stress GRNs genes, which are also thought to be linked to TEs, as reported by several studies suggesting that transposition events may be triggered during plant stress responses including salt [41], wounding [42], bacteria [43], and viruses [44]. We focused on the genes expressed in various stress conditions described by Barah et al. [45]. We searched in the 500 bp immediately upstream from these genes for Duster-specific, Brassicaceae-specific, and TAIR10-specific TEs (table 3). We found no enrichment of these upstream regions in Duster-specific TE copies (chi-squared *p*-value=0.73) or TAIR10-specific TE copies (chi-squared p-value=0.67). However, we found an enrichment for Brassicaceae-specific TEs (chi-squared p-value=5.1 × 10^−7^)

### TEs and transcription factor binding sites

The conservation and overrepresentation described above suggest a probably functional role for these TEs. We then investigated their ability to regulate gene expression, by assessing their ability to bind TFs. We investigated the co-occurrence of the transcription factor binding sites (TFBSs) identified with 27 TFs in ChIP-seq experiments by Heyndrick et al. [19] and the various TE annotations studied here.

We found that TFBSs were more frequently present in Duster regions (29.0%) than in Brassicaceae and TAIR10 regions (19.5%, and 14.6%, respectively, chi-squared *p*-values all < 10^−300^; Table 4). This pattern was even more marked for the analysis of method-specific regions: 53.9%, 29.7% and 24.9% of these regions, respectively, overlapped with TFBSs (chi-squared *p*-values all < 10^−300^). This trend was even stronger for analyses limited to the 500 bp immediately upstream from genes (49.3% for Duster-specific and 38.2% for Brassicaceae-specific TEs, chi-squared *p*-value < 10^−300^, TAIR10-specific TEs being untestable due to the low counts). Interestingly, these regions contained 567 Duster-specific and 48 Brassicaceae-specific TFBS regions, associated with more than seven TFs, and referred to hereafter as *hot TFBS*s. The identification of these regions suggests that there may be a hub of target genes involved in the important function of crosstalk between different processes [46].

**Table 4:** Method-specific TEs overlap with TFBS identified by ChIPseq. Occurrences are given between parenthesis.

|  | Hot TFBSs | TFBSs |
| --- | --- | --- |
| **All Duster** | 1.10% (1155) | 29.0% (30400) |
| **All Brassicaceae** | 0.51% (426) | 19.5% (16206) |
| **All TAIR10 TEs** | 0.40% (120) | 14.6% (4331) |
| **Duster-specific** | 2.89% (567) | 53.9% (10575) |
| **Brassicaceae-specific** | 0.93% (48) | 29.7% (1528) |
| **TAIR10-specific** | 1.13% (2) | 24.9% (44) |
| **5' Duster-specific** | 2.38% (114) | 49.3% (2365) |
| **5' Brassicaceae-specific** | 0.79% (7) | 38.2% (337) |
| **5' TAIR10-specific** | 3.33% (1) | 6.67% (2) |

An analysis of the 5’ sequences of flowering genes identified 6 key TFs known to be involved in flowering that could bind to both Duster-specific and Brassicaceae-specific regions (AGL-15, AP1, PI, AP3, SEP3, SOC1). Few sites for TFs involved in circadian rhythm and light response (PIF3, PRR5, PRR7) and development (GL3) were identified. Most were found in Duster-specific regions, with very few in Brassicaceae-specific regions, and none in TAIR10-specific regions. Interestingly, some Duster-specific regions were associated with several TFBSs.

We found 1757 and 1009 Duster-specific regions overlapping with crucifer and rosid CNS, respectively, and a TFBS. We found that 84 of these regions were highly conserved, as they overlapped with CNSs present in the 12 rosid species used for their identification, suggesting a presence in the common ancestors of the rosids more than 100 My ago. We also found that 9 of these highly conserved Duster-specific regions overlapped with a *hot TFBS*, suggesting the presence of a highly conserved hub of target genes involved in crosstalk between different processes. The top five highly conserved TFBSs from Duster-specific regions were AGL-15, AP1, SEP3, PRR5, and PIF4 (31, 23, 20, 14, 14 occurrences, respectively), all but one of which are directly involved in flowering process, the exception being PRR5, which is more closely related to circadian rhythms and light responses.

The CNSs associated with Duster-specific copies in the 500 bp upstream of the gene present in the 12 rosid species included 58 Duster-specific sequences: 42 target genes of floral regulators according to Chen *et al*. [40], 16 for which Duster-specific regions were colocalized with a highly conserved CNS and a TFBS (Table 5).

**Table 5:**
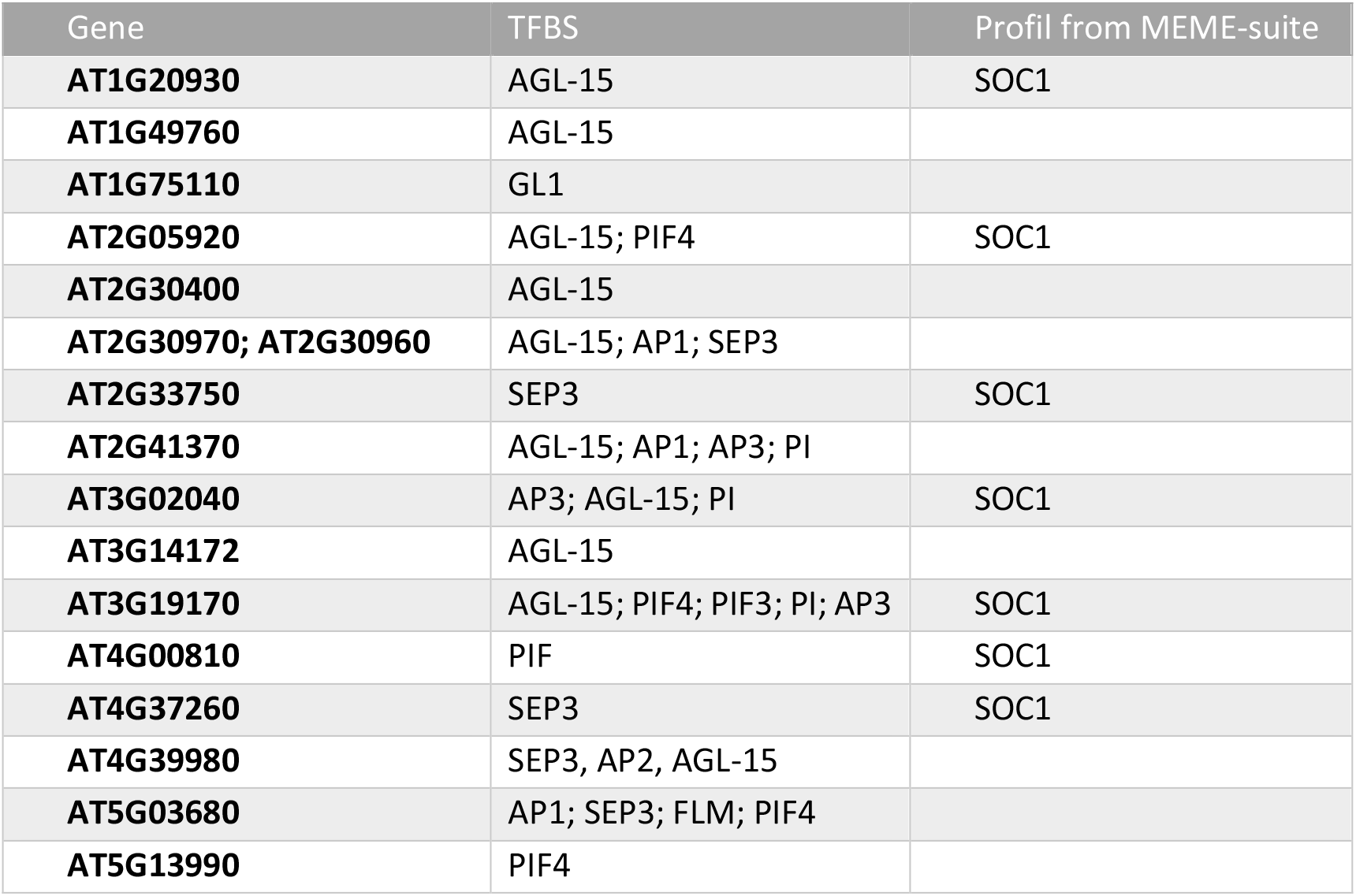
Sixteen floral regulators genes from Chen et al. [40] with a Duster-specific region in the 500 bp immediately upstream, colocalizing with highly conserved CNSs and TFBSs

Using the *MEME Suite*, we identified 7 of these genes as containing a sequence corresponding to a binding motif of the SOC1 TF, a MADS box factor active in flowering time control which may integrate signals from the photoperiod, vernalization and autonomous floral induction pathways. Thirteen of downstream genes were controlled by the floral regulation motifs of one or several type II TF-MADSs (AG, AP1, AP2, AP3, BLR, ETT, FLM, JAG, LFY, PI, RGA, SEP3, SVP).

## Discussion

### The need for a new dedicated repeat detection algorithm

RepeatMasker [47], Censor [48,49], and Blaster [19] are the tools most frequently used to annotate TE sequences in genomes. All these tools combine BLAST (or BLAST-like with seed and extend algorithms) calls with pre- and post-processing for the analysis of genomic sequences. They are all, therefore, subject to the intrinsic limitations of these algorithms, including a reliance on seeds to find alignments. These seeds in BLAST are *k-*mers with a default size of 11 nucleotides. BLAST requires two *k*-mers on the same diagnonal (i.e. alignment without gaps) to proceed with the analysis of an alignment to assess its relevance. An alignment score threshold determined with a probabilistic model is used for this assessment of relevance. These features may account for the lower sensitivity for this method than for Duster.

First, two *k*-mers are required to initiate an alignment. With the default BLAST parameters, this requires an exact match of at least 22 nucleotides between two sequences. This requirement can be decreased, as seed length is a parameter of BLAST that can be set, and it is decreased to 14 nucleotides for some implementations (seed size of 7 with WU-BLAST), but it still needs an exact match. For Duster, we allow mismatches in the *k-*mers, and the two k-mers may overlap. With the settings used for this analysis, we required a match of 21 nucleotides, but with some mismatches allowed.

Second, in the statistical test based on an alignment score threshold, even if the required exact match of 22 nucleotides is found, a gap-free alignment is produced for testing with the probabilistic model. The result depends on sequence length and on a model that is mathematically sophisticated, but too simple biologically, in that it considers successive nucleotides to be independent and equally probable. We now know that neither of these assumptions hold true for real sequences. Consequently, the model is of debatable value and may reject some alignments differently according to sequence lengths. In Duster, we retain all regions that match two *k-*mers, and the empirically chosen parameters yielded very few false positives (0.001).

We can see here that BLAST is not the most appropriate algorithm for finding small degenerate TEs. It was developed for a different purpose: identification of the best match within databases to a sequence used as a query. Its use to identify TEs constitutes a major deviation from its initial purpose, for which it performs well.

Duster was designed for the express purpose of finding old and degenerate TE copies. In addition to having a different *k-*mer strategy, it is essentially an alignment-free algorithm. BLAST attempts many alignments before reporting a match. Duster does not really require an alignment, just boundary coordinates, accounting for the greater speed of this algorithm. Boundaries may be considered imprecise as they are based on *k-*mers and their precision is therefore limited by *k-*mer size and the shift in the coordinates of the *k*-mer on the genomic sequence. With the parameters used here, the precision is about seven nucleotides. We think that this is sufficient for the identification of regions, and it may not be appropriate to aim for greater precision in the identification of very old, degenerate TE copies.

Improving sensitivity is always at the cost of specificity which is particularly difficult to evaluate in the context of an uncomplete TEs identification. Many TEs remain to be discovered even in a well studied genome such as *A. thaliana*. False positive rate is consequently difficult to measure. We proposed here a proxy for this assessment using CDS (see Materials and Methods) allowing to compare Duster with BLAST on the same basis. The good sensitivity performance of Duster is partly due to the chosen balance with specificity. However, sensitivity has also a cost in term of speed, and we think that Duster algorithm improve mainly on this feature and open the route for future new more efficient algorithms. Improving the sensitivity for BLAST algorithm to reach this of Duster, would have made this work more difficult or even impossible for computing reasons.

The work presented here highlights the utility of specifically developed tools for addressing certain difficult biological questions. It highlights the need for a new generation of sequence-finding tools, tailored to the particular biological question posed and perhaps replacing BLAST with more adapted algorithms.

### Long-term impact of TE copies

TEs are important sources of variation on which selection can operate in the evolution of species. Many examples of TEs generating new phenotypes have been reported in plants [50]. This phenomenon is known as “domestication” when the TE sequences are retrained in new genes, or “co-option” when TE insertions affect existing genes. TE sequences that become functional in the host are conserved by selection, which can be recognized over long periods of time. Other TE copies devoid of function in the host are progressively removed from the genome through the accumulation of point mutations and deletions. However, gene regulatory regions accumulate point mutations and deletions at a slower rate than other non-coding regions, because of their function. Consequently, TE insertions in these regions, even if neutral, may be difficult to remove once established.

TFs control the transpositional activity of TEs by binding to them, but they have also been shown to bind TEs in regions not supposed to be transcribed. Are they remnant of old TFBS or illegitimate because of a particular base composition of the regions? Whatever is the answer, the corresponding TFBS are, therefore, widespread throughout the genome. In some cases, TEs from the same family may be inserted close to several genes. This may lead to nearby genes being regulated by the same TF, potentially leading to their evolution into gene regulation networks. Genome-wide assessment revealed that hundreds of TEs have been co-opted into the regulatory regions of mammalian genes [51,52]. TEs have also been involved in both the creation of new regulatory networks [53,54] and the rewiring of preexisting ones [55]. Such networks are observed, for example, for the DAYSLEEPER gene in *A. thaliana* [56]. This gene has features in common with hAT transposases, suggesting that it may have been domesticated as a new TF. In order to operate, transposases are able to recognize TE DNA motifs close to the sequence boundaries in order to cleave DNA and initiate the transposition process. This suggests that this domesticated transposase may have conserved this property, as TF also bind specifically DNA motifs. This binding may be functional as it leads to the domestication of the transposase. Overall, this suggests that TE sequences, dispersed throughout the genome, are targeted by DAYSLEEPER, and may regulate host genes. Another interesting example is provided by the retrotransposon ONSEN in *Arabidopsis* [57]. Thieme *et al*. [58] showed that, following the heat stress-dependent mobilization of ONSEN, the progenies of treated plants contain up to 75 new ONSEN insertions. Progenies with additional ONSEN copies display broad environment-dependent phenotypic diversity. This finding suggests that some of the new TE insertions affect the expression of genes in a temperature-dependent manner. It also suggests that TE sequences may have contributed to individual local adaptation through the mutations they induce during bursts of transposition. Some of these bursts of transposition may result from activation by environmental stresses, promoting environment-sensitive phenotypes.

However, little is known about the long-term impact of TEs. Our CNS data suggest that some of the TEs identified were inserted in their current positions more than 100 My ago, during the Cretaceous period. The most important evolutionary event during this period was, perhaps, the spread of flowering plants (Angiosperms) to colonize the entire planet. Flowering plants were particularly successful at colonizing new areas and replacing the older established flora, with which they competed. TEs undoubtedly played an important role in this process. Some of the TE insertions we detected may, indeed, have played this role. Duster-specific copies appear to be old, degenerate, short, and surprisingly close to genes, lying in the 5’ flanking sequences known to correspond to gene regulatory regions. Their maintenance specifically in these zones suggests that they supply the host with a function, probably in the regulation of the neighboring gene.

The Duster-specific TEs identified here may have played an important role in building new pathways allowing flowering plants to adapt to their environment. Indeed, we found that Duster-specific TE copies were overrepresented in the 5’ regions of genes of the GRN for flowering (table 3). A significant proportion of these copies overlapped with TFBSs known to bind TFs involved in the control of flowering. Moreover, the histone

H3K27me3 mark was identified predominantly in method-specific TEs (see figure 3). This histone mark has been reported to be associated with genes expressed at low levels of in a tissue-specific manner [32], such as those involved in flower development.

Our results suggest a possible link between the success of flowering plants during the Cretaceous period and the co-option of TEs in the flowering GRN. However, further analyses are required to demonstrate a causal role. This study is a first step in this direction, identifying previously unknown candidate TEs.

Flowering has been studied in considerable detail, generating a wealth of data. The data used here are, therefore, clearly biased towards flowering. However, other impacts on other GRNs may subsequently be discovered with our ancient TE annotation, as and when new data become available.

Interestingly, our results suggest that identifying very old TE copies could facilitate the identification of TE-based regulatory modules selected a long time ago. They support the detection of TFBS in ChipSeq experiment, but also suggest a TE-based origin for many TFBS.

## Conclusions

In this study, we investigated the contribution of TEs to the bulk genome of Arabidopsis over a timescale that remains inaccessible to other approaches, through the use of a new tool that we developed, called Duster. Duster uses a new efficient algorithm, which identified an additional 10% of nucleotides as belonging to TEs. We have, thus, dug deeper into the dark matter than previous studies, leading to the recognition of old, degenerate TE sequences undetectable with other methodologies.

This study delivers a key result, improving our understanding of plant evolution and plant adaptation, by providing clues for identifying ancient TE remnants in gene regulatory regions underlying potential regulation modules. Some of the TE copies identified here may have been selected a long time ago, to drive adaption to changing environments.

## Data accessibility

The Brassicaceae TE copies from *Arabidopsis thaliana* Col-0 ecotype, *Arabidopsis lyrata, Capsella rubella, Arabis alpina, Brassica rapa* and *Schrenkiella parvula*, used for Duster annotation, and TAIR10 TE, Duster, and Brassicaceae annotations can be downloaded from https://urgi.versailles.inrae.fr/files/sequence/PubAthaDarkmatterData/PubAthaDarkmatterData.zip

The data are also available in a JBrowse genome browser at https://urgi.versailles.inrae.fr/jbrowse/gmod_jbrowse/?data=myData%2FAtha

## Supplementary material

The Duster code is available on github, and is distributed as part of the TEfinder package at https://github.com/urgi-anagen/TE_finder

## Acknowledgements

We thank Klaas Vandepoele for reading a first version of this manuscript and drawing our attention to interesting TFBS and CNS data that enabled us to improve the manuscript. We also thank Michaël Alaux and Johann Confais for their comments on the manuscript. This work was performed with the facilities of the Plant Bioinformatics Facilities (https://doi.org/10.15454/1.5572414581735654E12). Version 6 of this preprint has been peer-reviewed and recommended by Peer Community In Genomics (https://doi.org/10.24072/pci.genomics.100004)

## Conflict of interest disclosure

The authors of this preprint declare that they have no financial conflict of interest with the content of this article. Hadi Quesneville is one of the PCI Genomics recommenders.

## Notes

https://urgi.versailles.inrae.fr/files/sequence/PubAthaDarkmatterData/PubAthaDarkmatterData.zip

### Competing Interest Statement

The authors have declared no competing interest.

### Summary of Updates

Version 6 of this preprint has been reformatted according as recommended by Peer Community In Genomics

